# Evaluating bacteriophage impact on gut microbiome composition in broad-nosed pipefish (Syngnathus typhle)

**DOI:** 10.1101/2024.11.25.625175

**Authors:** Jamie Parker, Silke-Mareike Marten, Jelena Rajkov, Franziska I. Theising, Arseny Dubin, Olivia Roth

## Abstract

Bacteriophages play a crucial role in shaping microbial community dynamics in marine systems and have the potential to stimulate surges in pathogenic bacteria facilitating disease outbreaks. Notwithstanding, bacteriophages also serve as valuable biocontrol agents, underscoring their huge potential for aquaculture therapy treatments. Insights into the intricate tripartite interplay upon an infection with a virulent bacterium, its specific phages and the eukaryotic microbiome could improve our understanding of how bacteria-phage interactions behave in a natural microbial system. This investigation assessed the influence of a virulent *Vibrio alginolyticus* (K01M1) infection, in tandem with lytic (фSt2) and filamentous (фK04M1) *Vibrio*-specific phages, on the broad-nosed pipefish (*Syngnathus typhle*) gut microbiome using 16S rRNA amplicon sequencing and non-intrusive gastric swabbing. Microbial communities and diversity metrics were not permanently impacted by *Vibrio* and phage introductions, while the different infection regimes shaped *Vibrio*-specific dynamics. In the filamentous phage and *Vibrio*-only treatments, *V. alginolyticus* abundances spiked 12 hours post-ingestion. In contrast, *V. alginolyticus* numbers in the lytic phage and control treatment were significantly reduced, suggesting phage activity and specific elimination of the introduced bacteria. Assisted by relative true-gut tissue samples, a newly implemented non-intrusive swabbing method was successful at discerning the activity of two contrasting phages and supports previous work that encourages the use of фSt2 in bacteriophage treatments. Identifying *Vibrio*-specific phages with similar positive characteristics could be beneficial for the aquaculture trade, which is currently heavily impacted by the antibiotic crisis.

## 1. INTRODUCTION

Marine microorganisms have evolved along a continuum of symbiotic, commensal, opportunistic, and pathogenic states, with many capable of shifting activities in response to environmental changes (Wilson & Hastings, 1998; Thompson *et al*., 2004). In turn, changing environments often dictate the co-evolutionary outcomes of interacting species that exist within their realm of influence (Hendry & Kinnison, 1999; Reusch & Wood, 2007). Changes in the environment can influence pathogen fitness traits as indicated by increased replication rate (Mostowy & Engelstädter 2011; Wolinska & King 2009), contribute to immunologically compromised hosts (Wendling *et al*., 2013; Wendling & Wegner, 2013), and consequently impact virulence evolution (Anderson & May, 1982; Bull, 1994; Ewald, 1994; Ebert & Herre, 1996; Frank, 1996; Alizon *et al*., 2009; Engering *et al*., 2013). Biotic factors such as temperate bacteriophages can also contribute to increased pathogen virulence by supplementing bacteria with accessory genes (Waldor & Mekalanos, 1996; Wagner & Waldor, 2002; Austin *et al*., 2003; Payne *et al*., 2004). In contrast, some ‘phages’ have been shown to combat disease by lysing virulent disease agents, lending to the growing evidence that phage activities play an important role in manipulating the microbial environment and influencing the complex tripartite interactions between the eukaryotic host, bacteria, and phage (Levin & Bull, 2004; Rohwer & Thurber, 2009; Harrison & Brockhurst, 2017; Breitbart *et al*., 2018).

Phage lifecycles have evolved towards a number of subtle but distinguishable forms, each possessing unique characteristics and cellular repercussions. Lytic phages transmit horizontally by invading targeted bacteria, carrying out rapid phage replication and virion release following cell lysis, from which this lifecycle earns its name (Popescu *et al*., 2021; Chevallereau *et al*., 2022). Conversely, lysogenic phages are vertically transmitted without destroying the host cell, gradually integrating into the bacterial genome to form a ‘prophage’, which is then replicated following bacterial cell division (Weinbauer, 2004). Temperate phages are capable of switching between these two lifecycles, while filamentous phages can release virions without cell lysis in a chronic process or adopt a lysogenic state if needs dictate (Rakonjac *et al*., 2011; Sausset *et al*., 2020).

Lytic phages attracted interest for their therapeutic potential in the early 20th century (Summers, 2012; Wittebole *et al*., 2014), but the rise of antibiotics largely replaced phage therapy in the West. However, phage therapy has remained prominent in the East, especially in aquaculture, and is now being reconsidered in the West as an alternative to antibiotics (Nakai *et al*., 1999; Sulakvelidze *et al*., 2001; Nakai & Park, 2002; Merril *et al*., 2003; Defoirdt *et al*., 2011; Ryan *et al*., 2011; Oliveira *et al*., 2012; Richards, 2014). Bacteriophage therapy, as it is known, represents an alternative treatment against harmful ‘superbug’ bacteria that develop resistance to multiple antibiotics (Matsuzaki *et al*., 2005; Kortright *et al*., 2019). Identifying highly specific phage-bacteria interrelationships is often challenging; however, phage specificity can afford an advantage over non-specific antibiotic treatments with fewer unwanted side effects to the detriment of the host microbiome (Nieth *et al*., 2015). Despite its medical potential, bacteriophage therapy is still comparatively under-researched and would benefit from further systematic and fundamental *in vivo* studies that focus on pertinent virulent bacteria and their phage occupants.

*Vibrio* bacteria are the most abundant and diverse opportunistic bacteria in the marine realm, with some pathogenic phylotypes known to cause vibriosis in the natural environment (Blake *et al*., 1979; Thompson *et al*., 2004). These cases are often linked with seasonal salinity and temperature changes, with increased prevalence occurring during the warmer months (Martin *et al*., 2002; Dayma *et al*., 2015). Vibriosis is one of the most prevalent diseases affecting a wide range of crustacean, fish, and shellfish aquaculture practices, leading to significant economic losses and health concerns. Additionally, human exposure to contaminated aquatic environments, through ingestion or wounds, can result in severe infections (Baker-Austin *et al*., 2018; Ina-Salwany *et al*., 2019; Sanches-Fernandes *et al*., 2022).

Bacteria-phage studies are often carried out in the laboratory with *in vitro* liquid cultures with a single bacterial strain (Smith & Huggins, 1983; Pavlova *et al*., 1997; Gupta & Prasad, 2011; Cieplak *et al*., 2018). In addition, *in vivo* mammalian studies have used bacteriophage cocktails to assess the influence on bacterial targets, the immune system, and host-gut microbial communities (Maura *et al*., 2012; Febvre *et al*., 2019; Hsu *et al*., 2019). In fish, some aquaculture-related experimental studies on the gut have explored the phage protective qualities (Jun *et al*., 2013; Christiansen *et al*., 2014; Wendling *et al*., 2017; Cafora *et al*., 2019), but few assessed the phage-bacteria interactions on the microbiome (Donati *et al*., 2022). All these aforementioned studies used either extracted tissue or collected fecal samples to derive microbial assessments, raising questions regarding the potential for an alternative method that could be less intrusive and at risk of contamination. From an experimental perspective, there also appears to be a clear absence of comparative research assessing the effects of distinct bacteriophages adopting alternate lifecycles in parallel within the fish gut. A better understanding of individual phage influences on the tripartite relationship in the fish gut could aid in designing optimal phage cocktails for targeting specific bacterial diseases in farmed fish, while also providing insights into dynamics that may lead to crashes in wild fish stocks.

The broad-nosed pipefish (*Syngnathus typhle*) is part of the enigmatic syngnathid fish group, renowned for their bizarre morphologies and their unique male pregnancy evolution (Stölting & Wilson, 2007). Syngnathids have attracted interest due to their unusual immune repertoires, while other studies have advocated their use in phage-bacteria research (Wendling *et al*., 2017; Goehlich *et al*., 2019; Chibani & Hertel *et al*., 2020; Chibani & Roth *et al*., 2020; Roth *et al*., 2020; Parker et al., 2023). To investigate the influence of phage-bacteria interactions on the fish gut microbial composition over time, *S. typhle* was experimentally inoculated with either lytic or filamentous phages in combination with a known specific target, *Vibrio alginolyticus*. Repeated gastric swabbing and terminal gut tissue sampling in combination with 16S rRNA amplicon sequencing were used to assess the influence of the respective bacteriophages on the pipefish microbiome over time.

In line with previous syngnathid studies that were successful with *Vibrio* gut introductions, we hypothesized that **(i)** the introduced *V. alginolyticus* (K01M1) strain would be detectable in treatments from six hours post-ingestion onwards using nonintrusive gastric swabs. Successful phage therapy practices require high phage-bacteria interaction specificity. Previous reports suggest the phages utilized in this study have a strong specific affinity for *Vibrio alginolyticus*; therefore, it was hypothesized that **(ii)** all phage introduction treatments would not negatively impact the relative abundance of other non-*Vibrio* bacterial strains within the microbiome. Based on the physiological characteristics of the two bacteriophages used in this investigation, we hypothesized that **(iii)** treatments with lytic phage introduction would result in a reduced *V. alginolyticus* relative abundance in comparison to all other treatments, while due to a less destructive lifecycle, **(iv)** filamentous phages would exhibit similar *V. alginolyticus* abundances to phage-devoid treatments.

## 2. MATERIAL and METHODS

### 2.1 Ethics statement

All research conducted in this investigation is in accordance with German animal welfare law and the ethical approval provided by the Ministerium für Energiewende, Landwirtschaft, Umwelt, Natur und Ditgitalisierung (MELUND) Schleswig-Holstein under the permit number: Ves.-Nr.: V 242-35168/2018 (63-7/18). All fish used in this study were aquarium-bred, and the species is not endangered.

### 2.2 The eukaryotic host

*Syngnathus typhle* originating from wild-caught populations of the Baltic Sea were bred for several generations and reared at GEOMAR aquaria facilities in Kiel at 18 °C. Before the experiments, fish were fed twice a day with a mixture of frozen and live mysids.

### 2.3 The bacteria

*Vibrio alginolyticus* is a prevalent gram-negative bacterium commonly found in the Baltic Sea and other marine domains (Oberbeckmann *et al*., 2011; Böer *et al*., 2012). *V. alginolyticus* (K01M1) samples were revived from cryopreserved cultures previously isolated from nine healthy *S. typhle* individuals (Roth *et al*., 2012; Wendling *et al*., 2017). The K01M1 strain was selected for this investigation based on previous reports and laboratory trials confirming its selective vulnerability to infection by the two phage sub-types used here (Wendling *et al*., 2017).

Isolate subsamples were preserved in 15 PSU liquid Medium101 (0.5% (w/v) peptone, 0.3% (w/v) meat extract, and 1.5% (w/v) NaCl and 25% glycerol in Milli-Q water) and thawed gradually at 25 °C while shaken at 180 rpm overnight. A day later, samples were diluted (1:10) in fresh medium and grown for a further 1.5 h (22 °C at 180 rpm) to encourage exponential culture growth. Samples were subsequently centrifuged (8000 rpm at 12 °C), supernatant was removed, and pellets were washed in 40 mL sterile seawater. Wash steps were repeated twice more prior to pellet resuspension and sample pooling in a total of 45 ml. Two dilution series (−1 to −7) were created in phosphate buffer saline (PBS), and 100 µl was incubated on 101_15PSU agar plates (22 °C) for 16 h. Plate count numbers were as follows: 7×10^7^ (MW cfu/mL) and 3.5×10^7^ (cfu/500 µL), which was used for the the inoculum in this experiment.

### 2.4 The bacteriophages

One lytic and one filamentous bacteriophage were selected for this experiment based on their proven ability to specifically infect the K01M1 strain, as demonstrated in laboratory trials and a previous study (Wendling *et al*., 2017). The lytic phage (фSt2) was originally isolated from the coastal waters of Crete in Greece (Kalatzis *et al*., 2016) and obtained in liquid culture (2×10^9^ mL^−1^) from the University of Copenhagen, Denmark. To prepare фSt2 samples, 2 x 1600 µL aliquots were centrifuged for 2 min (13,000 rpm), supernatants were filtered through 0.2 µm filters, and products were stored in Medium 101 in the fridge at 4 °C. *V. alginolyticus* K01M1 overnight cultures were diluted (1:100) and incubated for 1.5 h at 22 °C and consistently shaken at 180 rpm. For phage re-cultivation, 4.5 mL of bacterial culture was mixed with 500 µL of фSt2 and incubated for a further 4 h at 22 °C while being shaken at 180 rpm. The supernatant was collected following 2 min centrifugation (13000 rpm) then filtered again through a 0.2 µm filter and stored at 4 °C. The final фSt2 concentration used for the experiment was 3×10^9^ pfu/mL.

The filamentous phage (фK04M1) was previously extracted from a *V. alginolyticus* strain of the same name (Wendling *et al*., 2017). A subsample of cryopreserved K04M1*Vibrio alginolyticus* bacteria culture in 15 PSU liquid Medium101 was thawed overnight at 25 °C while being shaken at 180 rpm. After 26 h, bacteria were pelleted following centrifugation at 13000 rpm and the supernatant was fed through a 0.2 µm filter to isolate the фK04M1 phage culture. A final concentration of 5×10^10^ pfu/mL was used for фK04M1 during the inoculation experiment.

### 2.5 Spot assays

In line with a previous study, standard spot assays were carried out for both phage types to confirm phage activity on a plate overlaid with Baltic seawater with 0.4 % agar and 200 µL with the susceptible K01M1 culture (Wendling *et al*., 2017). Plates were kept overnight (~20 h) before assessing the effectiveness of each phage by the presence (susceptible) or absence (resistant) of plaque formation.

### 2.6 Experimental design

To understand the influence of lytic and filamentous phage cultures on the gut microbiota of *S. typhle*, a time series experiment was carried out that assessed how colonization with *Vibrio* bacteria and bacteriophages, respectively, changed the host microbiomes over a period of 16 days. Live mysid cultures were incubated for 30 min with either *V. alginolyticus* K01M1 only (**K1**) (500 µl - K01M1 and 500 µl - Medium101 (15 PSU)), lytic phage фSt2 and K01M1 (**LY**) (500 µl - фSt2 and 500 µl - K01M1), filamentous phage фK04M1 and K01M1 (**FI**) (500 µl - фK04M1 and 500 µl - K01M1) or sterile seawater (**CO**) (500 µl - Medium101 and 500 µl - seawater) (Fig. 1a) (supplementary; Table. S4).

**Figure 1.**
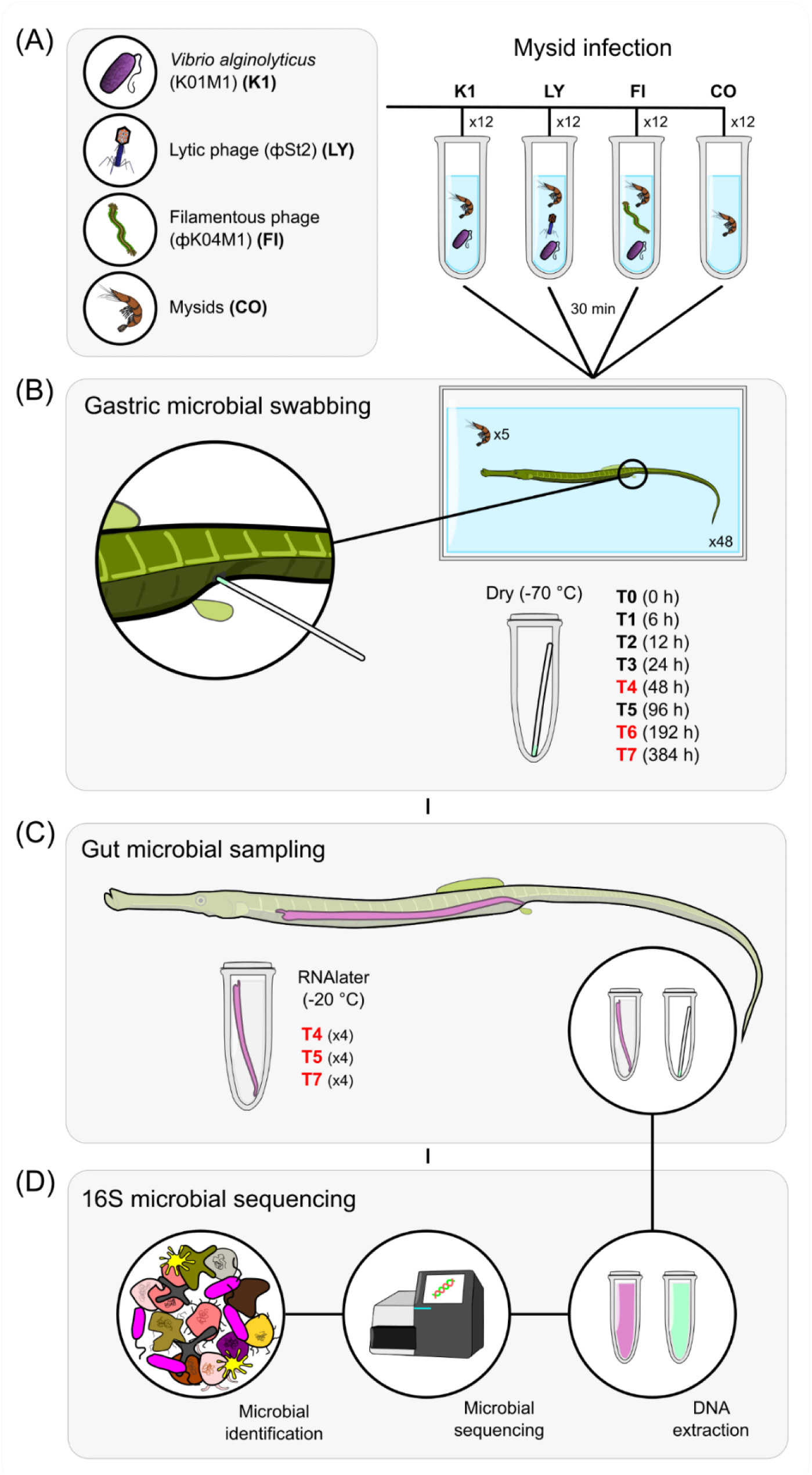
Refined schematic diagram depicting the following experimental steps used during this investigation: (A) Mysid incubation with bacteria and phage mixtures; (B) gastric microbial swabbing over a 16-day period; (C) gut microbial sampling at three latter stages of the experiment; and (D) 16S amplicon sequencing and microbiome assessment.

Fish were placed in individual 20 L tanks and allowed to acclimate for one week in addition to being starved for 48 h prior to the experiment. Five treated mysids were added to each tank for consumption, with 12 fish replicates used per treatment group initially. To sample the microbial communities occupying the end of the pipefish digestive tract, sterile absorbent paper points (29 mm) (Antæos) were used to swab the gastric pore/terminal gut opening of each fish (Fig. 1b). Care was taken when handling fish out of the aquaria, and the sampling area was meticulously dried and disinfected using ethanol prior to swabbing to minimize environmental contamination. Gastric swabbing was carried out prior to mysid consumption (T0), and then after 6 h (T1), 12 h (T2), 24 h (T3), 48 h (T4), 96 h (T5), 192 h (T6), and 384 h (T7). For timepoints T0-T4, 12 swab replicates were used, while at timepoints T5, T6, and T7, there were 8, 8, and 4 swab replicates, respectively, per treatment. This replicate reduction was due to fish being euthanized for gut tissue sampling. All swabs were stored dry at −70 °C in preparation for DNA extraction.

Gut tissue samples were dissected from four fish per treatment at time-points T4, T6, and T7, resulting in a total of 12 fish per treatment over this period (Fig. 1c). Consequently, the total number of swab samples at T5, T6, and T7 was reduced to 32, 32, and 16, respectively. Fish were euthanized with a lethal dose of MS-222 (Tricaine, 500 mg L^−1^; Sigma-Aldrich, Munich, Germany) before gut dissection, gut content removal, and gentle washing with PBS. Samples were initially placed in RNAlater at 4 °C before relocation to −20 °C until DNA extraction. To minimize the chances of contamination, all gut dissections were carried out under the sterile bench.

### 2.7 DNA extraction, PCR amplification and sequencing

Thawed microbial swab tips and gut samples were homogenized using 180 μL of lysozyme solution (20 mM Tris CL pH 8.0, 2 mM sodium EDTA, 1.2% Triton X-100, lysozyme 20 mg/ml) in lysing matrix A-tubes (MP Biomedicals) using a tissue shredder (Qiagen). For DNA extraction, the DNeasy PowerSoil Kit and standard protocol were used (Qiagen, Hilden, Germany), in addition to an improved Gram-positive bacteria pre-treatment used previously (Korsch *et al*., 2018; Tanger *et al*., 2024). DNA quality was then assessed using a NanoDrop-1000 spectrophotometer (NanoDrop), and amplification of the V3-V4 hypervariable 16S rRNA gene region was targeted (341F-805R). Library preparation was carried out at the Institute for Experimental Medicine (UKSH, Kiel), and paired-end sequencing (2 × 300 bp) was conducted via Illumina MiSeq (Illumina, USA) at the Institute of Clinical Molecular Biology (IKMB) in Kiel (Fig. 1d).

### 2.8 Data analysis

Amplicon read sequences were demultiplexed before primer cutting, filtering, chimera removal, sequence quality checks, and denoising using DADA2 via QIIME 2 (v2022.8). By utilizing the V3-V4 region as a reference, the Silva (v132) (Quast *et al*., 2012) was used for ribosomal rRNA alignment and taxonomic assignment. Chloroplast and mitochondrial-associated 16S rRNA sequences were removed before continuing with downstream analyses.

Initial alpha rarefaction curves were used to confirm the sampling success of microbial communities and the observable clarification of species at various sequencing depths. Resulting Operational Taxonomic Units (OTUs) were carried forward for analysis using Phyloseq (v1.42) (McMurdie & Holmes, 2013) in R (v4.2.2) (R Core Team, 2013). Sequence variants without a clear taxonomic class were removed prior to consolidating variants into ASV-level groups. Taxa present in fewer than four samples and with less than two counts were filtered prior to converting the remaining taxa to relative abundance levels.

#### 2.8.1 Diversity analysis

Local mean species diversity (α-diversity) among microbiome strain constituents was assessed using Faith’s phylogenetic diversity (Faith PD) and Shannon diversity and observable genera measures via the microbiome package (Lemos *et al*., 2011; Lahti & Shetty, 2017). Analysis of variance (ANOVA) and post-hoc Tukey-testing were carried out to assess significant differences between specific treatments and timepoints.

To assess β-diversity across all treatment samples, a Bray-Curtis dissimilarity matrix was utilized for permutational multivariate analysis of variance (PERMANOVA) (999 permutations). This was carried out on a two-factorial blocked PERMANOVA (data ~ treatment + timepoint) and repeated measures PERMANOVA (data ~ treatment * timepoint). Further PERMANOVAs were conducted on treatments within each individual timepoint as well as pairwise PERMANOVAs conducted on a *Vibrio* data subset (999 permutations).

To assess the effectiveness of the gastric swabbing technique at determining accurate *Vibrio* community proportions, relative abundance swab data were correlated with the matching gut tissue samples extracted at timepoints T4, T6, and T7.

#### 2.8.2 Principal component analyses

Principle component analysis (PCA) was carried out on each timepoint subset to visualize treatment differences. Statistical assessments across PC1-4 were conducted using multivariate analysis of variance (MANOVA) for each timepoint representation. If significant differences were identified (MANOVA; *p* < 0.05), individual ANOVAs were used to identify the most influential principal components, on which post-hoc Tukey-tests were performed to extract treatment differences.

To assess changes in *Vibrio* populations across treatments, an exclusive *Vibrio* data subset was established, excluding all taxa not aligning with the *Vibrio* genus. The same PCA and statistical tests conducted on the full taxa datasets were carried out. *Vibrio* strain loading values were imprinted upon PCA visualizations that highlighted significant treatment disparities in order to help with the interpretation of treatment-specific influences of each *Vibrio* strain. *Vibrio* strain deductions were based on NCBI nucleotide BLASTN+ reports (v2.16.0) (Altschul *et al*., 1997; Camacho *et al*., 2009), with the top 100 hits considered. As *V. alginolyticus* was introduced in this study, efforts were made to identify units with a high likelihood of a positive *V. alginolyticus* match. In turn, units with > 5 *V. alginolyticus* hits, 100% query coverage, and the highest max score were interpreted as *V. alginolyticus*. To support strain similarity deductions, a phylogenetic tree of the *Vibrio* DNA sequences was constructed using the TN93 (Tamura-Nei, 93) model. *V. alginolyticus* combined relative abundance fluctuations were also assessed throughout the experimental period, with ANOVA and post hoc Tukey’s tests being used to determine significant differences between treatment and timepoint combinations.

## 3. RESULTS

### 3.3 Microbial diversity

To assess the general localized microbial diversity in this investigation, α-diversity measures were adopted (Faith PD, Shannon diversity, and observable species). Significant Faith PD-derived phylogenetic differences were observed between swab treatments (*P* = 0.037) and timepoints (*P* = 4.6×10^−12^) in isolation but not in the interaction between the two (supplementary; Table S8). In terms of treatment effects, Faith PD was found to be significantly greater in FI compared with CO (*P* = 0.03). T0 was shown to have a significantly higher Faith PD than four other timepoints (T2, T3, T4 and T7), while T2 exhibited a significantly reduced diversity (T0, T1, T5 and T6). Three timepoints (T2, T3, and T7) showed a significantly lower Faith PD compared to T6, while three other timepoints (T0, T1, and T6) showed a significantly higher Faith PD compared to T3 (supplementary; Table S9).

Similar microbial diversity trends could be observed in the Shannon diversity, with ANOVA results for Shannon diversity differences highlighting significant differences between treatments (*P* = 0.0014) and timepoints (*P* = 3.22×10^−8^), but not the interaction between the two (supplementary; Table S8). Post hoc Tukey’s tests uncovered a significantly greater Shannon diversity in the K1 treatment compared with both CO (*P* = 0.008) and LY (*P* = 0.007). In regard to specific timepoint differences, Shannon-diversities of T0, T2, and T7 were all shown to be significantly reduced to those of T4, T6, while T5 was shown to have a significantly higher Shannon-diversity than to five different timepoints (T0, T1, T2, T3, and T7) (supplementary; Table S10). As with the other α-diversity assessments, treatment (*P* = 0.018) and timepoint (*P* = 8.57×10^−9^) were shown to be significantly different when assessing observable species, while the interaction was not (supplementary; Table S8). In line with the Faith PD assessments, the only significant treatment effect was between FI and CO, with FI exhibiting a higher number of observed species (*P* = 0.02). Post hoc testing revealed that T2 had significantly fewer observed species compared to five other timepoints (T0, T1, T4, T5, and T6), while T3 showed significantly fewer species than in three timepoints (T1, T5, and T6). T5 had a significantly greater number of observed species than four other timepoints (T1, T2, T3 and T7), while T6 had a greater number than three other timepoints (T2, T3, and T7) (supplementary; Table S11).

Gut sample Faith PD and observed species α-diversity analyses showed there to be no significant treatment or timepoint effect across timepoints T4, T6, and T7 (supplementary; Fig. S2; Table. S16). Shannon diversity analysis of the gut sample data set only showed a significant difference between timepoints, with post hoc testing showing (*P* = 0.01).

For gastric swab samples, comparative microbial β-diversity (Bray-Curtis) assessments between treatments and timepoints were conducted. Non-metric multidimensional scaling (NMDS) of the Bray-Curtis dissimilarity matrix for all relative abundance swab data exhibited a robust overlap of all treatments and disparities with the water and swab controls (supplementary; Fig. S3). After removing water and swab controls, β-diversity assessments utilizing constructed Bray-Curtis dissimilarity matrices were explored (supplementary; Fig. S4). PERMANOVA assessments showed significant differences for the main factors, treatment and timepoint, in isolation (*P* = 0.001), while the interaction between treatment and time was not found to be significant (supplementary; Table. S19). Focus was given to each treatment’s microbial community within timepoint subsets, as this provided the optimum comparative view into the microbial states for each treatment and their progression compared to the others of that time. Following PERMANOVA, no differences were observed between treatments from within each timepoint data subset (supplementary; Table. S19). For the gut samples, a significant difference was found between timepoints; however, this was not the case for treatment and the treatment-timepoint interaction (supplementary; Table. S22).

Swab relative abundance observations showed a *Vibrionaceae* spike at 12 hours (T2) in each treatment (supplementary; Fig. S5). Further examination at the genus level highlights a large portion of these taxa as *Aliivibrio*, while a smaller subset represents *Vibrio*. Two factorial repeated measures PERMANOVA showed a significant treatment and timepoint effect, while also showing a trend in the interaction which encouraged further pairwise comparison assessments. *Vibrio* β-diversity (Bray-Curtis) was found to be significantly different in FI and K1, when compared to CO at timepoints T1 (FI: *P* = 0.001, K1: *P* = 0.001), T2 (FI: *P* = 0.003, K1: *P* = 0.001), and T4 (FI: *P* = 0.007, K1: *P* = 0.001). K1 *Vibrio* diversity was also found to be significantly different than in LY at T1 (*P* = 0.008), T2 (*P* = 0.001), and T4 (*P* = 0.03), while at a reduced significance, FI was found to be different to LY at T5 (*P* = 0.043). Lastly, at T2, CO *Vibrio* diversity was found to differ from that of LY (*P* = 0.036) (supplementary; Table. S20). In terms of comparing *Vibrio* relative abundances across treatments and timepoints using ANOVA, the only significant difference was found between treatments (*P* = 0.03) (Fig. 2; supplementary; Table. S21).

**Figure 2.**
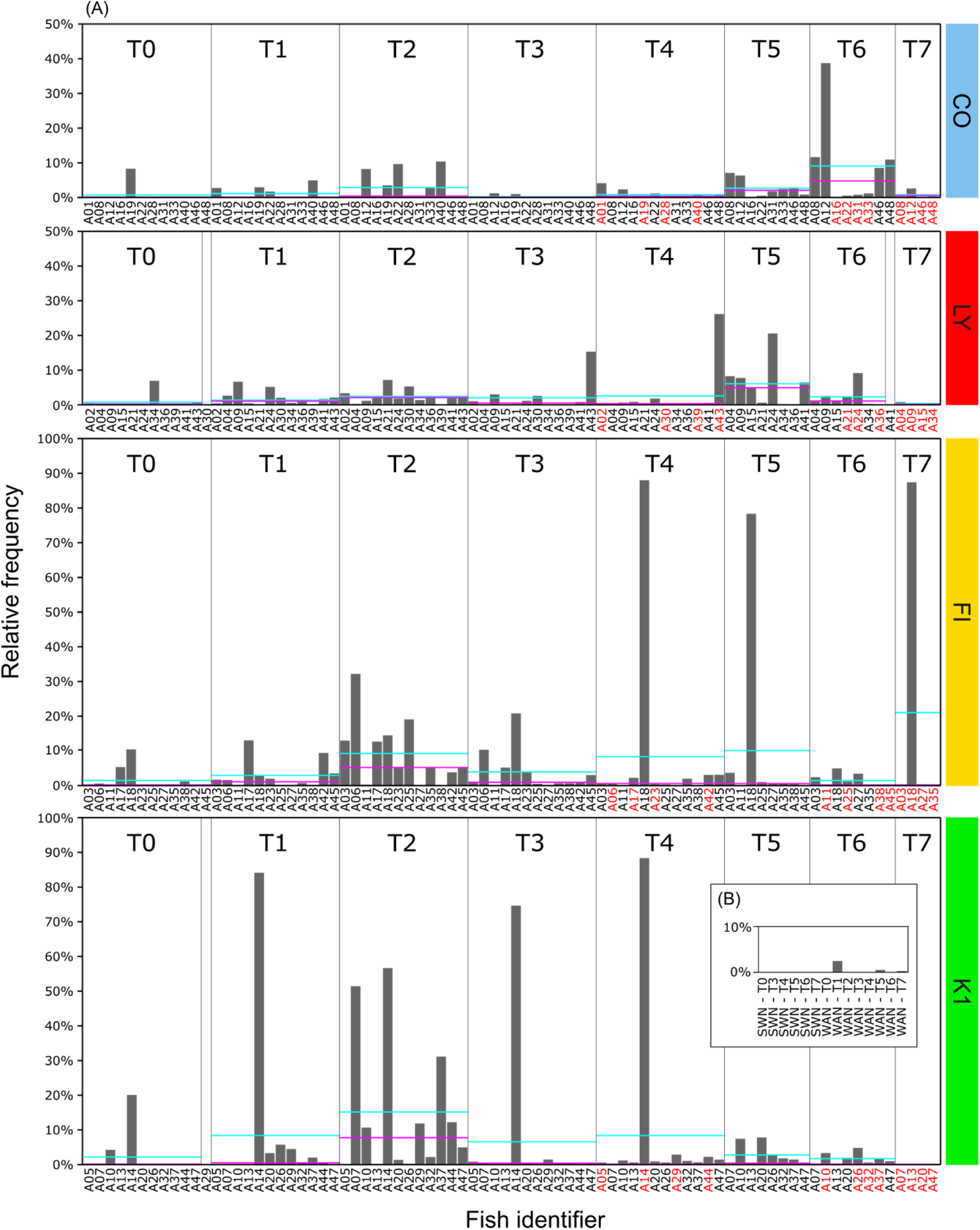
(A) *Vibrio* relative abundances for each timepoint in control (CO), lytic phage (LY), filamentous phage (FI), and *V. alginolyticus* (K1) gastric swab samples. (B) Swab (SWN) and water (WAN) negative controls indicated in cyan and pink lines indicate abundance mean and median values, respectively. Fish identifiers are derived from the unique resident tank identifier, and those highlighted in red at T4, T6, and T7 indicate fish used for gut tissue samples.

Insufficient replicates with *Vibrio* populations rendered comparative statistical analysis between treatments unfeasible for gut samples (Fig. 3). However, a positive correlation was found between *Vibrio* relative abundances from gastric swabs and matching gut replicates (R2 = 0.39, *P* = 1.6×10^−6^) (supplementary; Fig. S7).

**Figure 3.**
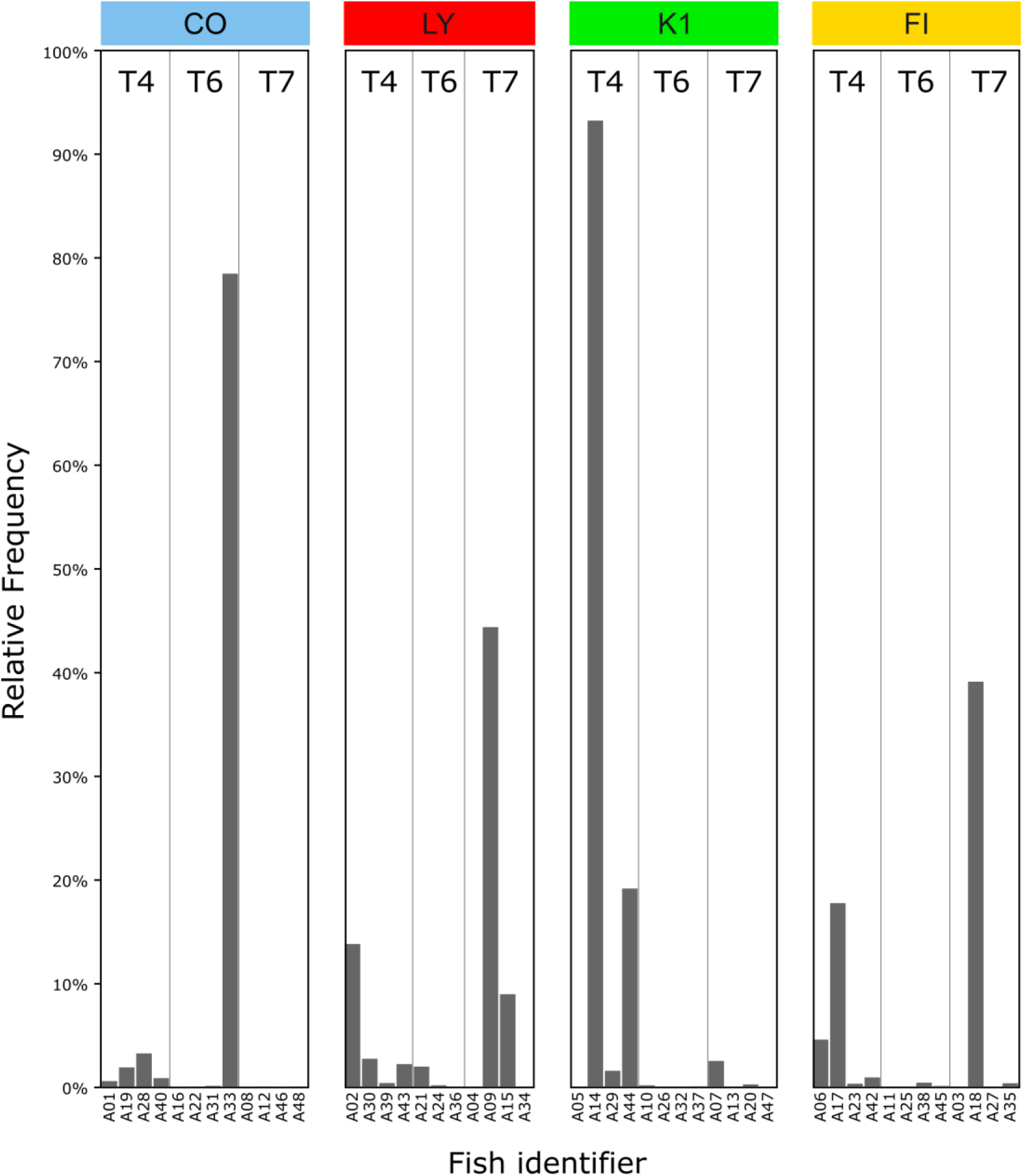
(A) *Vibrio* relative abundances for each timepoint in control (CO), lytic phage (LY), filamentous phage (FI), and *V. alginolyticus* (K1) gut tissue samples. Fish identifiers are derived from the unique resident tank identifier.

### 3.4 Principal component analysis

Whole microbiome timepoint PCA plots (PC1:4) displayed a consistent sample overlap, with MANOVA assessments indicating no significant differences between the treatments across PC1 to PC4 (supplementary; Fig. S8 and Table. S24). The only significant difference observed when carrying out principal component ANOVA for each timepoint was PC1 of T3 (*P* = 0.03). In this instance, PC1 explained differences in the microbial communities of FI and LY replicates (supplementary; Table. 24).

Exclusive *Vibrio* PCA plots for each timepoint explain more variation with the formation of clusters evident at T2 in particular (Fig. 4) (supplementary; Fig. S9). MANOVA assessments for each timepoint (PC1:4) showed significant treatment differences at T1 (*P* = 0.005), T2 (*P* = 0.0002), T4 (*P* = 0.049), and T5 (*P* = 0.04) (supplementary; Table. S26). ANOVA assessments on T1 principal component scores showed PC1 and PC2 to harbour significant treatment differences (*P* = 0.02 and 0.03, respectively); post hoc Tukey tests indicated that the FI and CO pairwise comparison was significantly different (PC1) (supplementary; Table. S26). Treatment differences were most distinct at T2, with PC1 explaining the majority of the difference following ANOVA testing (*P* = 1.1×10^−8^). Post hoc Tukey’s tests further highlighted the existence of significant differences between the control (CO) and each of the other three treatments, FI (*P* = 1×10^−6^), K1 (*P* = 1.1×10^−8^), and LY (*P* = 0.003). Additionally, LY *Vibrio* communities were shown to be significantly different from K1 (*P* = 5.1×10^−5^) and FI (*P* = 0.005) (supplementary; Table S26).

**Figure 4.**
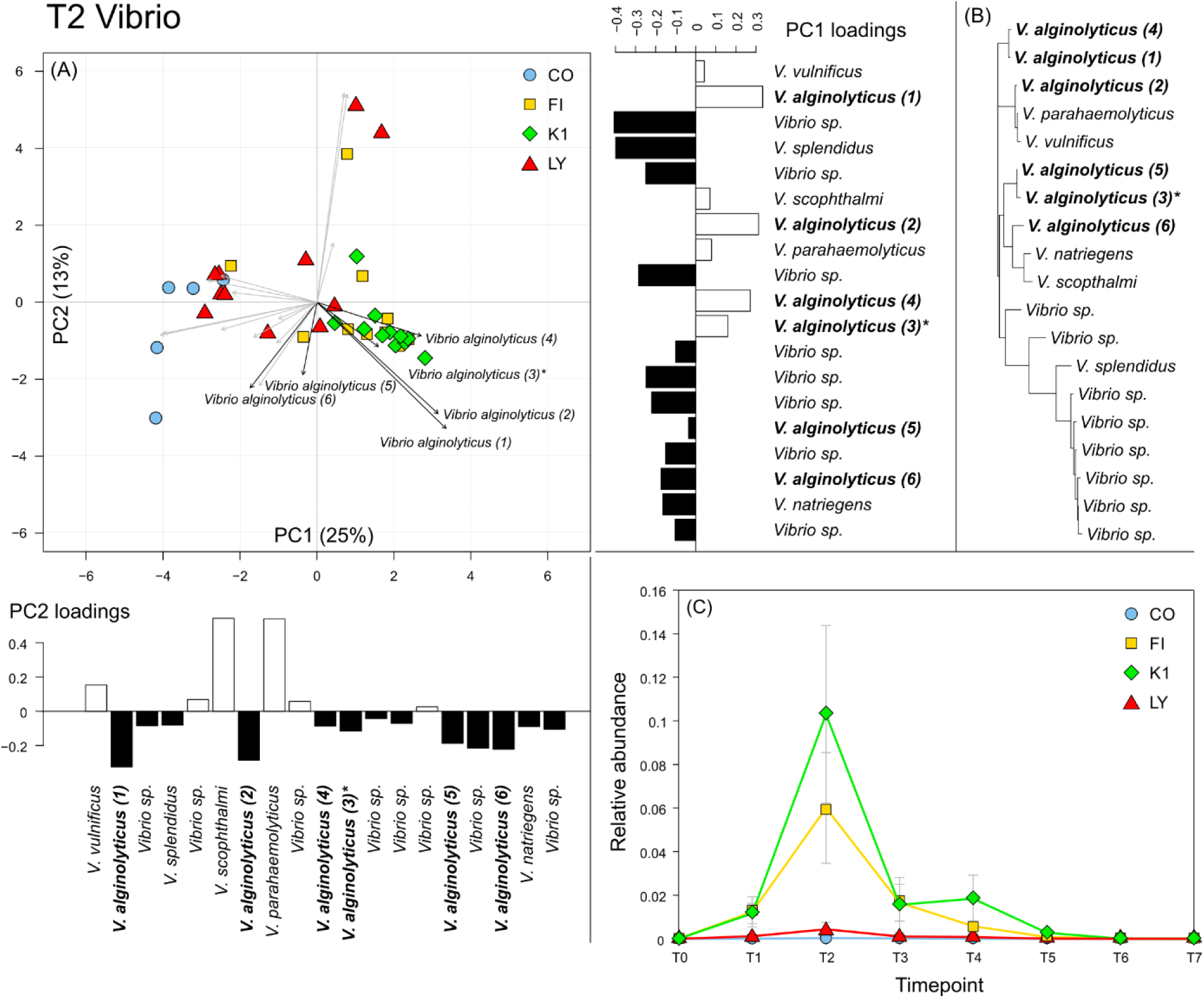
(A) PCA component plot of *Vibrio* strain data subset at T2 (12 hours), with control (CO), filamentous phage (FI), *Vibrio* (K1), and lytic phage (LY) treatments. Loadings for all *Vibrio* strains influencing the spread of replicates are represented by dispersing arrows, with dark labeled arrows indicating *V. alginolyticus* strains and the introduced K01M1 strain (*). *Vibrio* strain loading values for PC1 and PC2 are also represented in the flanking bar charts, with positive and negative loadings signified in white and black, respectively. (B) Phylogenetic tree based on *Vibrio* strains influencing treatment differences at T2, created utilizing the TN93 (Tamura-Nei, 93) model. (C) Combined relative abundance of all *V. alginolyticus* over the course of the experiment.

*Vibrio* PCA loadings at T2 show nineteen *Vibrio* strains with different modes of influence of the four treatments (Fig. 4 and supplementary; Fig. S5). Eight of these taxonomic units were identified to genus level (*Vibrio* sp.), and in the context of PC1 loadings, they were all positively associated with LY and CO replicates, but not FI and K1. Another six units were identified as potential *V. alginolyticus* strains, which is supported by phylogenetic observations derived from a high sequence similarity among all six units (Fig. 4b). One sequence in particular (*V. alginolyticus* (3)) matched specifically with the introduced K01M1 strain used in this study. Taking into account the loading influence of the six *V. alginolyticus* strains for both PC1 and PC2, four units were positively associated with K1 and FI treatment replicates at T2. Combined *V. alginolyticus* relative abundances at each timepoint for each treatment showed a spike at T2, similar to that of the overall *Vibrio* assessments (Fig. 4c). At this timepoint, ANOVA an post hoc testing highlighted K1 *V. alginolyticus* abundances to be significantly greater than those in CO (*P* = 6.6×10^−9^) and LY (*P* = 3.3×10^−8^) (supplementary; Table. S27). *V. alginolyticus* abundances were also found to be significantly greater at T2 in FI replicates compared with CO (*P* = 0.02).

### 3.5 Gut samples

Gut samples taken at timepoints T4, T6, and T7 (n = 4) were intended to support microbial swab assessments. Gut timepoint PCA plots depicted a similar trend to swab communities, with all treatment replicates except some disparate K1 individuals at T4 (supplementary; Fig S10). No significant differences were found between treatments following MANOVA at timepoints T4, T6, and T7, in line with the corresponding swab timepoint analyses (supplementary; Table S25).

## DISCUSSION

The complex relationship that exists between phage, bacteria, and eukaryotic host, compared with dual interaction-based research, has expanded a breadth of new research topics, attempting to understand the potential threats, symbiotic benefits, and environmental influences that surround the tripartite interaction (Wendling *et al*., 2017; Marchi *et al*., 2023). Considering the world’s seas, understanding this relationship could be of particular importance, as temperature and salinity fluctuations are commonplace and have been shown to influence virulent bacterial blooms, impact aquaculture stocks, and highlight impending problems with the warming climate (Lipp *et al*., 2002; Kimes *et al*., 2012; Vezzulli *et al*., 2012; Baker-Austin *et al*., 2013; Archer *et al*., 2023). Research on marine fish has deconstructed the three-way relationship by assessing the phage-bacteria interactivity *in vitro* (Goehlich *et al*., 2021; Goehlich *et al*., 2024), while others have explored the dynamics of the triplicity through the lens of a eukaryotic host immune response and its benefits within the aquaculture trade (Nakai *et al*., 1999; Wendling *et al*., 2017). This investigation builds on these connections by exploring the impact on the eukaryotic host gut microbiome, a feature that has been comparatively under-researched (Donati *et al*., 2022), while adopting a continuous non-intrusive sampling method.

*S. typhle* gut microbial communities (16S rRNA) were assessed following the introduction of the opportunistic *V. alginolyticus* in tandem with *V. alginolyticus*-specific filamentous and lytic bacteriophages. The impact of the bacteriophage treatments on the eukaryotic host microbiome was investigated using comparative α- and β-diversity metrics. Significant differences in microbial diversity were observed within treatments and timepoints, respectively. Aside from a single T3 PCA result that displayed a weak distinction between FI and LY communities, comparisons between treatments within individual timepoints highlighted no significant differences, suggesting that phage/*Vibrio* treatments did not alter the fundamental eukaryotic host microbiome in this investigation. This is in line with several mammalian studies that have reported that high lytic phage doses do not stimulate a significant change in gut microbial diversity (Tanji *et al*., 2005; Golomidova *et al*., 2007; Mai *et al*., 2010; Grubb *et al*., 2020), while others emphasized that regular phage dosage can result in the opposite (Bao *et al*., 2018; Hsu *et al*., 2019; Lin *et al*., 2019). This observation in the context of phage therapy is a beneficial contrast to antibiotic use, which has been shown to disturb the host gut microbiome (Patangia *et al*., 2022; Fishbein *et al*., 2023).

In a force-fed salmon experiment that attempted to understand the passage time differences between dry and soaked food pellets, 50% of the gastric tract was found to be emptied between six and twelve hours post-feeding (Aas *et al*., 2017). Another salmon study used inert faeces markers to understand gastrointestinal evacuation times of different diets, finding that all three food remnants could be detected after 12-15 hours (Storebakken *et al*., 1999). Similar passage times could be attributed to *S. typhle* in this investigation, as a *Vibrio* spike was observed at T2 (12 hours), likely indicating the passing of the ingested *Vibrio*-laced food. *V. alginolyticus* in particular was highly represented at T2 in FI and K1 treatments, and the K01M1 strain was successfully identified, supporting the first hypothesis of this investigation (i). These results, in addition to positive correlation with gut tissue samples, endorse the use of gastric swabbing in this investigation as a cohesive, non-intrusive approach to gut microbial assessments. Swab results from T2 also stand as an important marker, signifying comprehensive gut exposure to the treatment with subsequent timepoints.

The lytic phage life cycle comprises bacterial infection, rapid intracellular replication, and host cell destruction (Sulakvelidze *et al*., 2001; Weinbauer, 2004; Abedon, 2008). A conclusive, modular, bacterial host-range framework for bacteriophages has not been forthcoming, but it is generally understood that most phages are capable of infecting within a single bacterial genus (Ackermann & DuBow, 1987; Weinbauer, 2004; Flores *et al*., 2011), with a few exceptions (Paolozzi & Ghelardini, 2006; Dekel-Bird *et al*., 2015; Walsh *et al*., 2023). Species range with a single genus can vary greatly among phages (Kauffman *et al*., 2018), but there is evidence to suggest that phages are more likely to infect a bacterium of the same clade from which it was derived (Wendling *et al*., 2018). Lytic phages (фSt2) used in this experiment appear to have been operating specifically on the *V. alginolyticus* that were introduced collectively, with reduced relative abundances in LY matching more with CO, which did not receive bacterial input. This could indicate that the phages introduced in the LY treatment have lysed a high proportion of *V. alginolyticus* bacteria, in contrast to FI and K1, where in the latter no phages were introduced, while in FI the filamentous phages rather resemble a chronic infection that does not lyse their bacterial hosts (Sausset *et al*., 2020). Strain specificity in the lytic фSt2 is also supported by the fact that total *Vibrio* relative abundances showed no significant difference between LY and the other treatments. Therefore, results suggest that an upturn in lytic activity and the converse reduction in targeted bacterial numbers by T2 look to have occurred, and LY replicates share a more similar growth pattern with CO, supporting the third hypothesis of this study (iii).

Filamentous phages operate at a slower pace than their lytic counterparts, gradually releasing intracellularly formed virions in a chronic process that can slow cell growth without inducing cell lysis (Sausset *et al*., 2020). PCA assessments at T2 demonstrated FI to be very similar to K1 replicates in terms of its *Vibrio* population structure, strongly influenced by *V. alginolyticus* units. Interestingly, *V. alginolyticus* numbers in FI were significantly higher than in CO but not when compared to LY relative abundances. This could be an indication that the filamentous phage used in this experiment is inhibiting *V. alginolyticus* cell growth. At T2, the perceived earliest point of detection, the *Vibrio*-only treatment (K1) exhibited a significantly higher relative abundance of *V. alginolyticus* compared to CO and LY. This is a coherent finding, as K1 replicates were free from destructive phage activity. Moreover, K01M1’s positive influence in distinguishing the FI/K1 pair from CO/LY highlights that incorporated FI phages are operating in a chronic form rather than lytically, strongly supporting the third hypothesis of this study (iv).

A previous study found that the фSt2 lytic phage used in this investigation was able to successfully infect a range of *V. alginolyticus* strains, as well as one *V*. *parahaemolyticus* and one *V*. *harveyi* strain, demonstrating a broad host range for *Vibrio* bacterial strains (Kalatzis *et al*., 2016). Similarly, the filamentous фK04M1 has been shown to be an effective infector of *V. alginolyticus*, having been extracted from the same bacterial strain used in this investigation (Skliros *et al*., 2016; Wendling *et al*., 2017). In this regard, фK04M1 phages have been shown to preferentially infect within the *V. alginolyticus* clade from which they were derived (Wendling *et al*., 2018). It is difficult to conclusively determine whether other bacterial strains were affected by the phage treatments used in this investigation, as minor fluctuations in smaller bacterial communities can be influenced by a number of other factors (Dong & Gupta, 2019; Barreto & Gordo, 2023). However, based on the α- and β-diversity assessments, as well as the lack of combined *Vibrio* abundance differences between treatments in this study, bacterial communities appear consistent in their fluctuations across all treatments. This indicates that single treatments likely did not impact the microbiome in the broadest sense and therefore support the second hypothesis of this study (ii). It should be noted that an alternative outcome may have been reached if multiple treatment doses had been implemented, a trend that has been discussed previously (Bao *et al*., 2018; Hsu *et al*., 2019; Lin *et al*., 2019). The insurance of an uncompromised host microbiome following phage introduction is a requisite foundation of an effective phage therapy treatment (Kutter *et al*., 2015). Phage therapy has been carried out successfully in a number of aquatic species, including salmon, oysters, and corals (Cohen *et al*., 2013; Higuera *et al*., 2013; Jun *et al*., 2014). In turn, the practice has huge potential, particularly for the aquaculture trade, which can rely too heavily on the use of antibiotic treatments (Romero *et al*., 2012). фSt2 has a number of related studies advocating its potential use in phage therapy treatments (Kalatzis *et al*., 2016; Skliros *et al*., 2016; Kalatzis *et al*., 2018). While multiple dosing assessments and further validation would be beneficial to clarify фSt2 specificity and identify potential harmful side-effects that can result from its use (Kalatzis *et al*., 2018), this investigation provides additional support for фSt2’s use as a *V. alginolyticus* clade-specific treatment.

This investigation raised a number of challenges and considerations that could be useful for future microbial experiments adopting a similar tripartite structure. The consistency of the microbial results accrued using the swabbing method and the disparities with water/swab negative controls suggest the method was successful, non-intrusive, and free from environmental contamination. It is possible that the series of swabs were in part hindered by the reduction in sample size due to gut tissue sampling at timepoints T4, T6, and T7, rendering the results more stochastic compared with the initial timepoints. Gut samples were chosen in this investigation as a comparative supplement; however, there is a lack of contextual input with regards to the earlier stages. For this strategy to be successful, future investigations should include gut samples at the earlier stages and in numbers that are not to the detriment of the swab replicates. On reflection, it stands to reason that retaining the swab replicate numbers throughout the experiment would have been more beneficial for understanding the latter gastric microbial communities. Based on the overall diversity fluctuations found across treatments in this experiment, the introduction of *V. alginolyticus*/phage cocktails did not evoke a detrimental or permanent change in the eukaryotic host gut microbiome. Previous studies have reported that repeated phage doses can stimulate compositional shifts in the gut microbiome of mammals (Bao *et al*., 2018), which could also be the case in this study if repeated feedings were carried out. Future investigations should explore this point, as it could have implications for applied phage therapy treatments.

This study provides useful insights into the nuances of the phage-bacteria-host tripartite interaction and highlights subtle differences in activity between lytic and filamentous phages in the host gut environment. The lytic phage findings in this study support previous affirmations for фSt2’s use in *Vibrio*-specific bacteriophage therapy, which could benefit industrial aquaculture practices as an alternative or supplement for antibiotic treatments in the future.

## Data Accessibility

Raw sequencing data is available in NCBI SRA under project PRJNA1186123 at www.ncbi.nlm.nih.gov/sra/PRJNA1186123.

## Funding

Funding provided by the German Research Foundation (DFG) (RO-4628/4-2; WE 5822/1-2; RO-4628/3-1), from the European Research Council (ERC) under the European Uniońs Horizon research and innovation program (MALEPREG: eu-repo/grantAgreement/EC/H2020/755659) to OR, and the Collaborative Research Center CRC 1182 Origin and Function of Metaorganism funded by the German Research Foundation (DFG). Computational work was carried out using the NEC HPC system of the (Christian-Albrechts-University) University Computing Centre in Kiel.

## Acknowledgments

We would like to thank all members of the Marine Evolutionary Ecology group and those at the GEOMAR institute for their assistance with aquaria maintenance and scientific discussion, namely: Freya Pappert, Kim-Sara Wagner, Ralf Schneider, Haqdil Shad, Johannes Hasse, and Fabian Wendt. We would also like to thank Panos Kalatzis for generously supplying the lytic bacteriophages used in this investigation and Carolin Wendling for her conceptual input. Additional thanks are given to the DFG and ERC for financing this project’s research.

## Author contributions

OR conceived the study. JR, SM, FT and OR conducted experiments and processed fish samples. JP, AD and OR analysed the data and wrote the manuscript.

## Notes

### Competing Interest Statement

The authors have declared no competing interest.

