## Supplemental figures for "Evaluating bacteriophage impact on gut microbiome composition in broad-nosed pipefish (Syngnathus typhle)"

### Supplementary figures

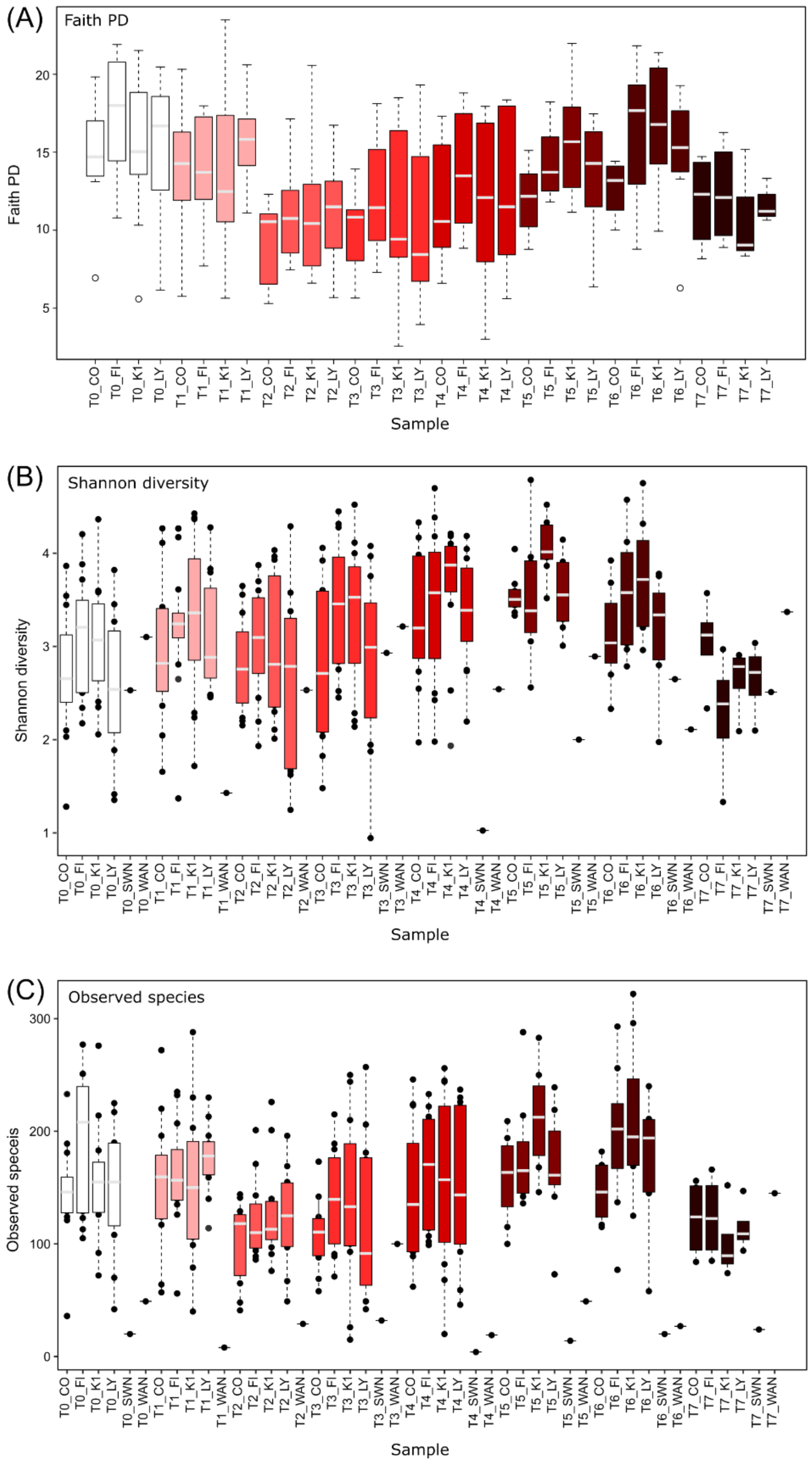


**Figure 1.** (A) Faith diversity, (B) Shannon diversity and (C) Observed species boxplots for swab samples depicting microbial diversity changes through time (T0-T7) for control (CO), filamentous phage (FI), Vibrio (K1) and lytic phage (LY) treatments. Negative swab (SWN) and water (WAN) samples also included in Shannon and Observed species representations.

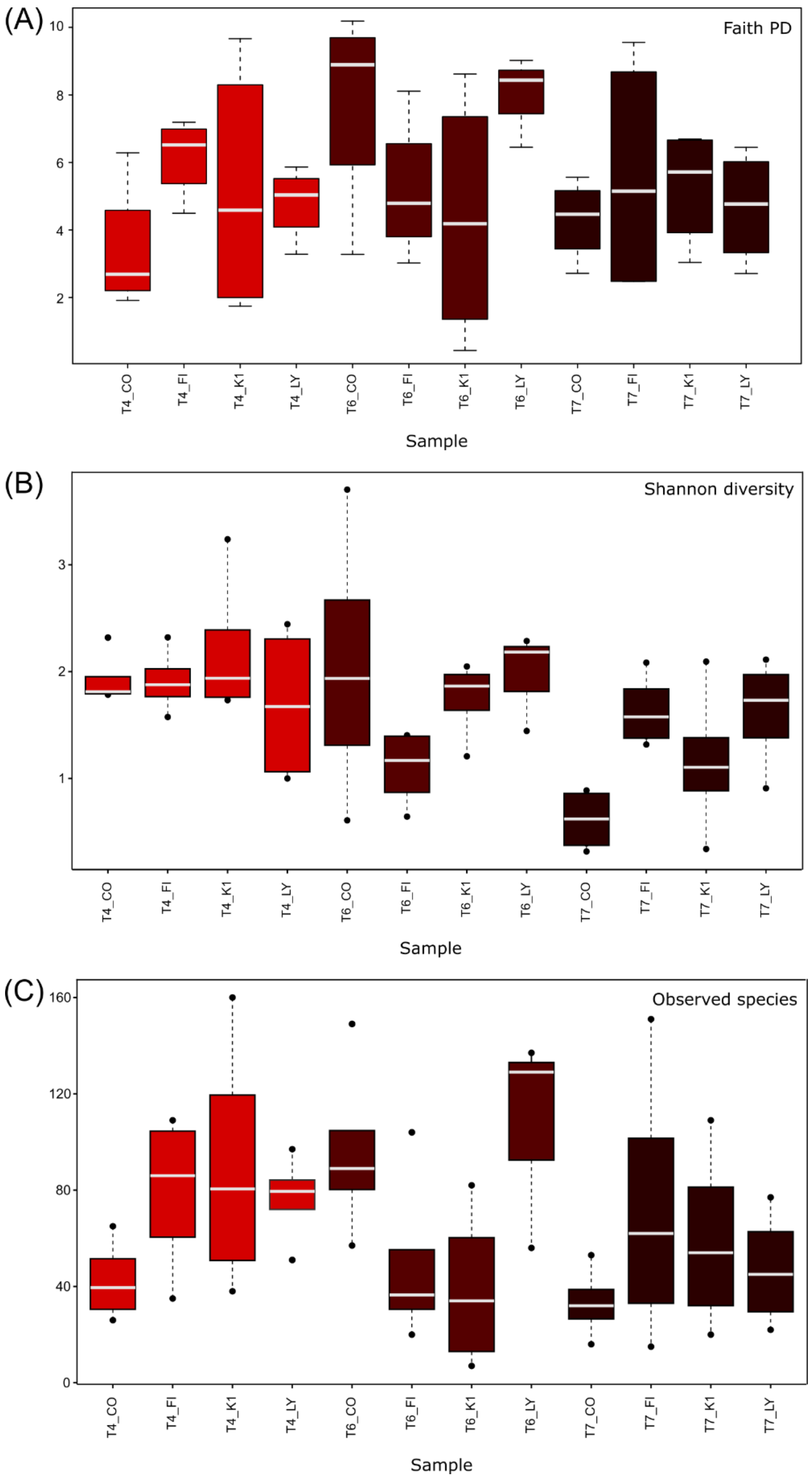


**Figure 2.** (A) Faith diversity, (B) Shannon diversity and (C) Observed species boxplots for gut samples depicting microbial diversity changes through time (T0-T7) for control (CO), filamentous phage (FI), Vibrio (K1) and lytic phage (LY) treatments.

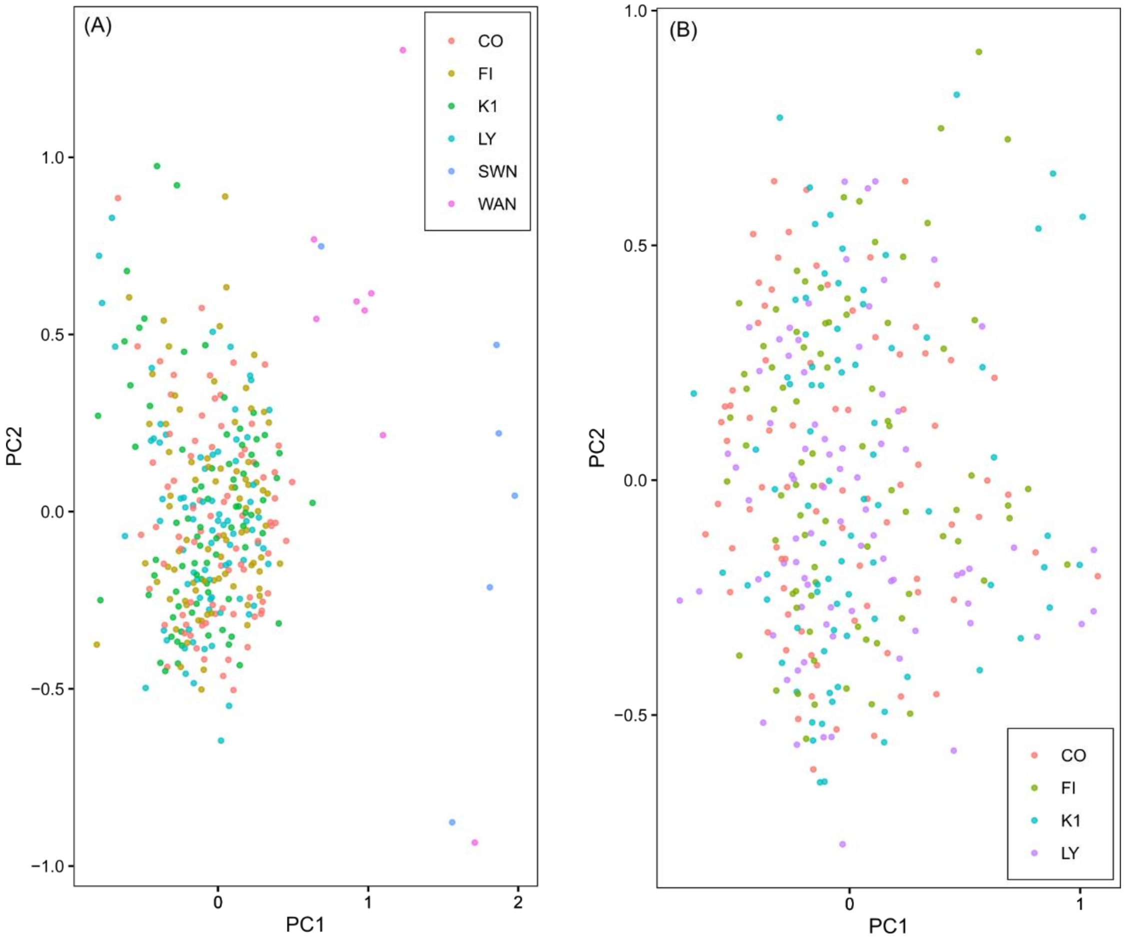


**Figure 3.** (A) Non-metric multidimensional scaling (NMDS) plot of control (CO), filamentous phage (FI), *Vibrio* (K1), lytic phage (LY) treatments, in addition to water (WAN) and swab (SWN) controls. (B) NMDS plot with all treatments, excluding water and swab controls.

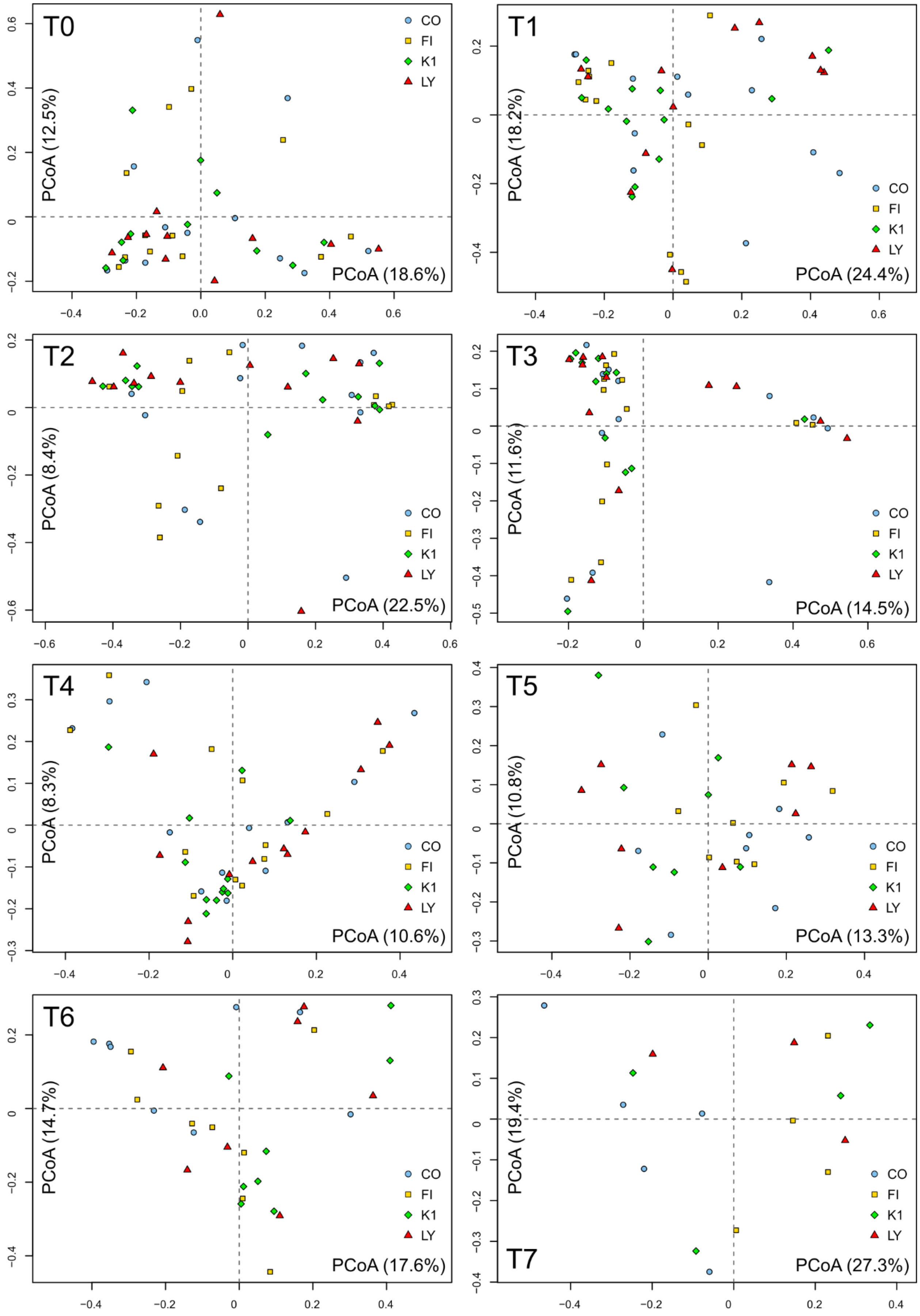


**Figure 4.** Principal coordinate analysis of relative abundances at each timepoint for all treatments: Control (CO), filamentous phage (FI), *Vibrio* (K1) and lytic phage (LY).

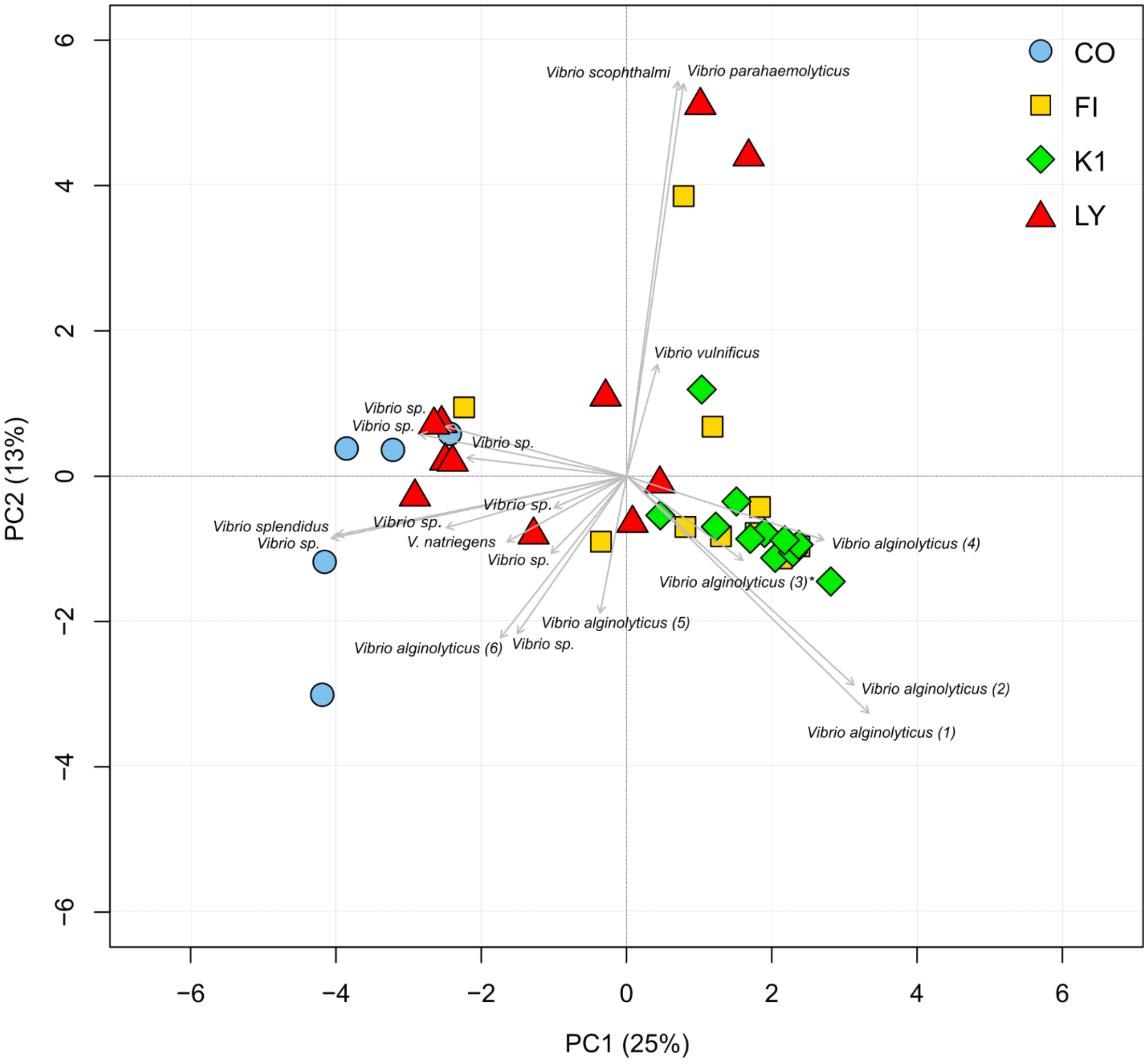


**Figure 5.** PCA component plot of *Vibrio* strain data subset at T2 (12 hours), with control (CO), filamentous phage (FI), *Vibrio* (K1) and lytic phage (LY) treatments. Loadings for all *Vibrio* strains influencing the spread of replicates are represented by dispersing arrows, and introduced (K01M1) strain (*).

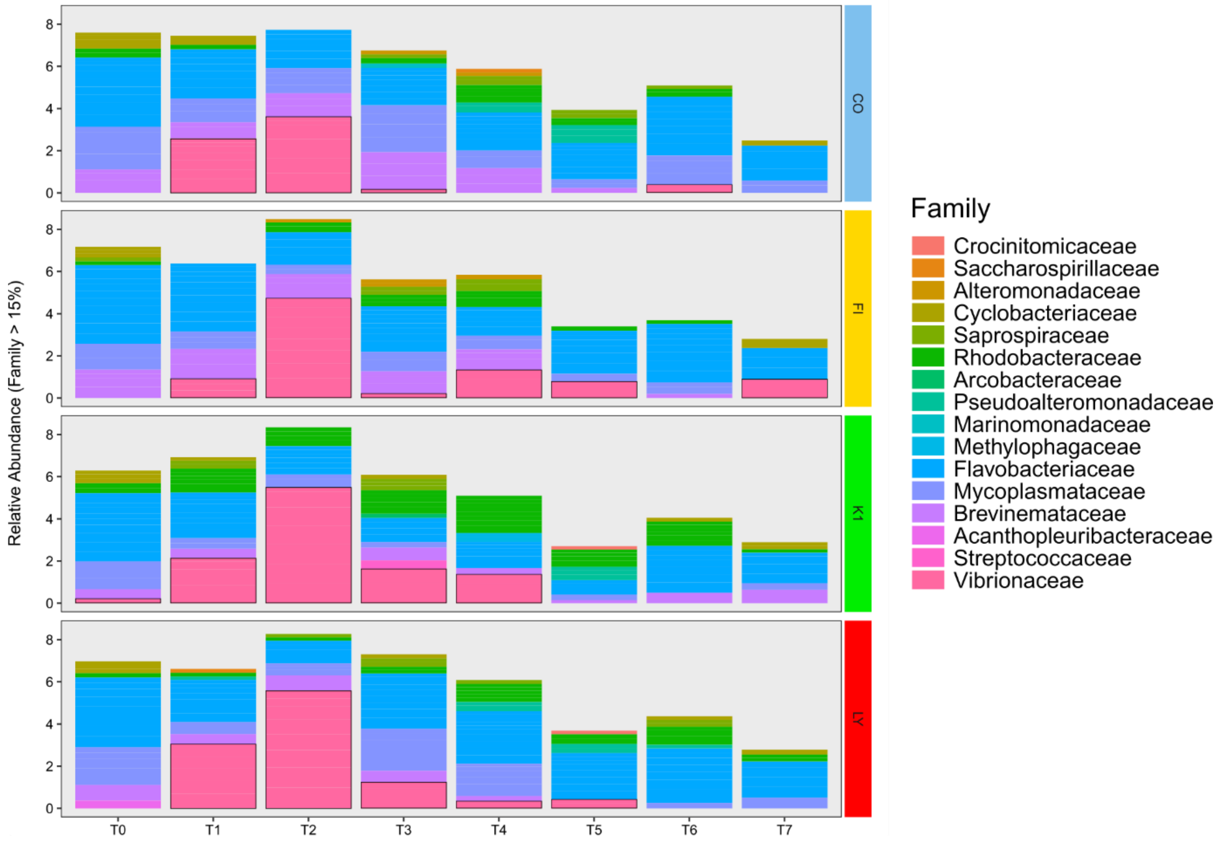


**Figure 6.** Stacked bar chart depicting relative abundances of microbial families (>15%) over each timepoint and for each treatment. Dark rimmed sections highlight Vibrionaceae.

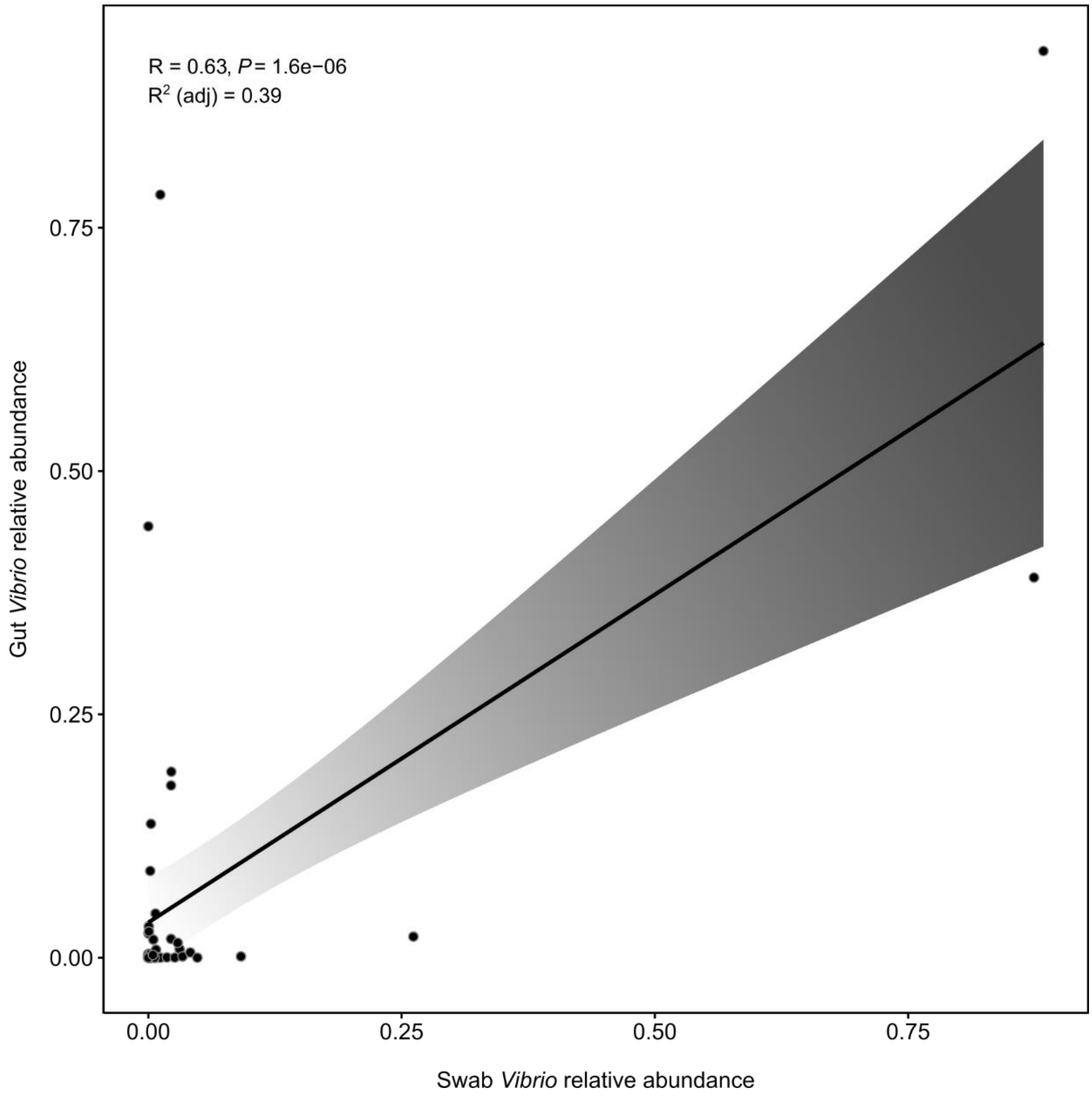


**Figure 7.** Pearson's correlation of between matching swab and gut replicates for all treatments at timepoints T4 , T6 and T7.

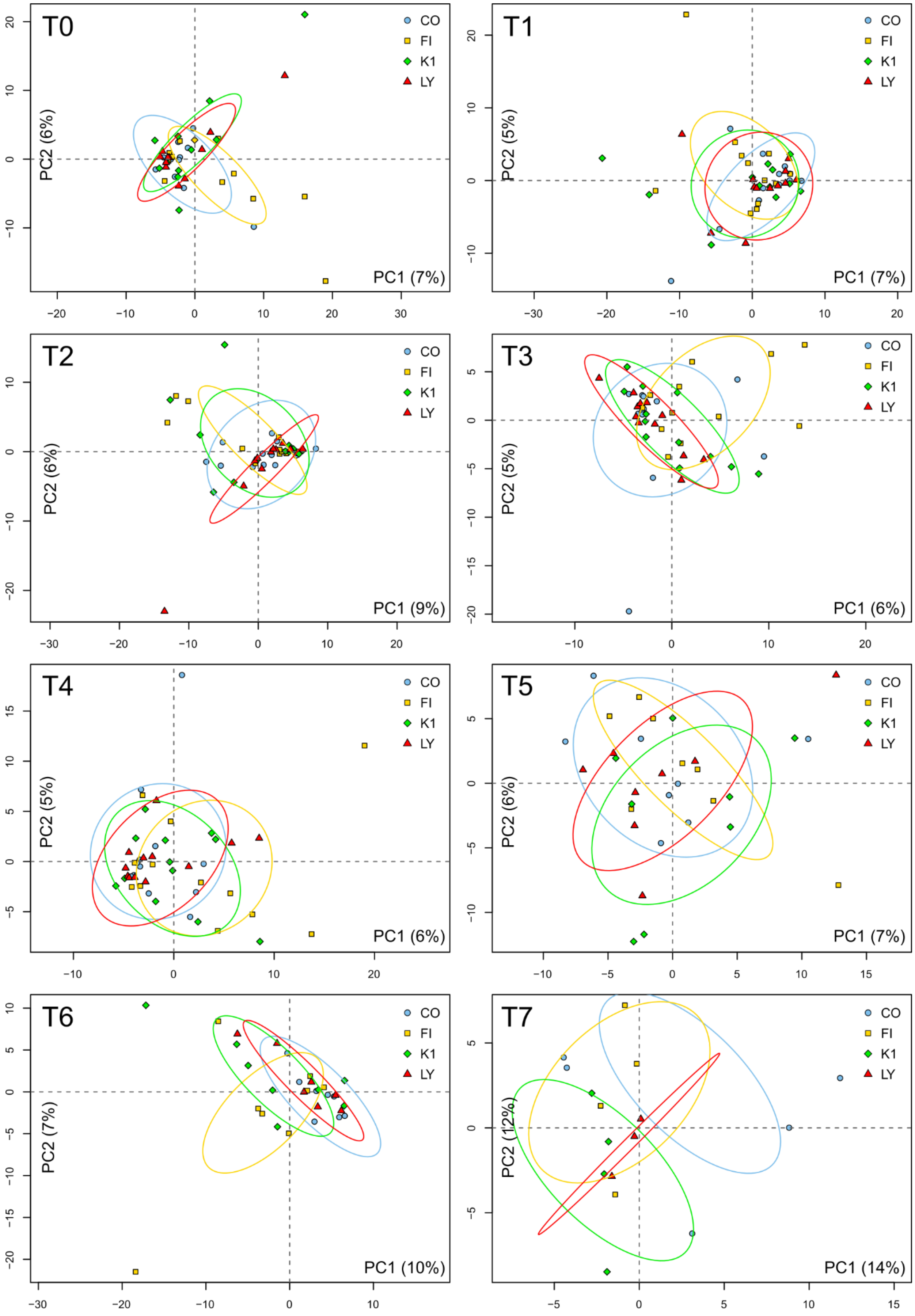


**Figure 8.** Principal component analysis plot of microbial relative abundances (swab) at each timepoint, and treatment: control (CO), filamentous phage (FI), *Vibrio* (K1) and lytic phage (LY).

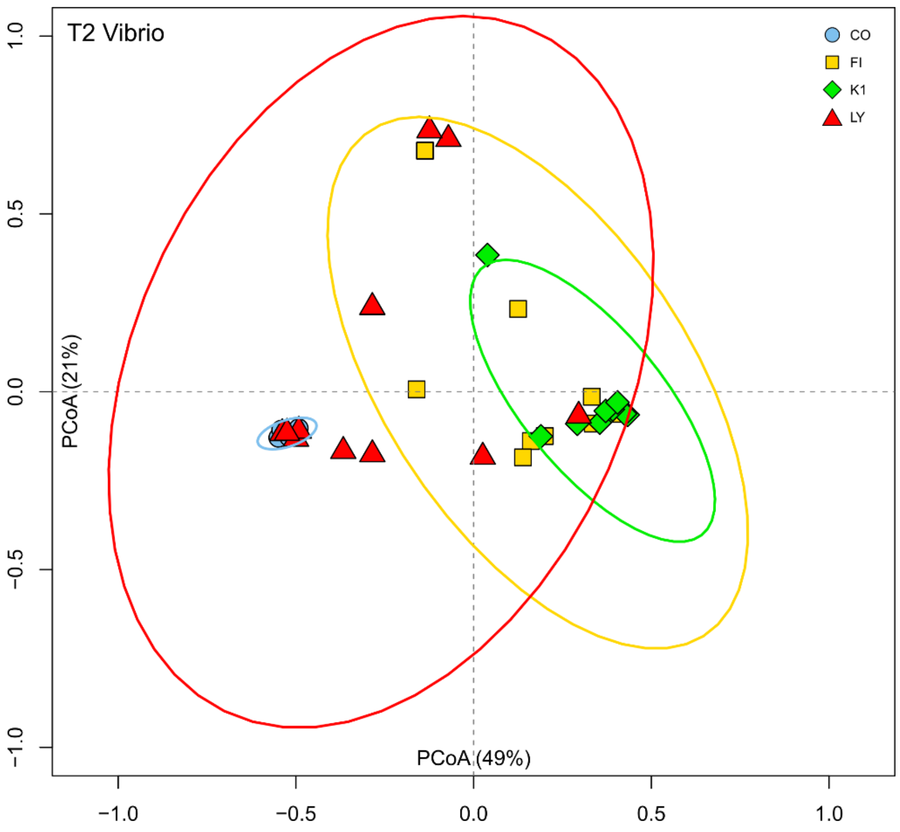


**Figure 9.** Principal coordinate analysis of *Vibrio* relative abundances at T2 for control (CO), filamentous phage (FI), *Vibrio* (K1) and lytic phage (LY) treatments.

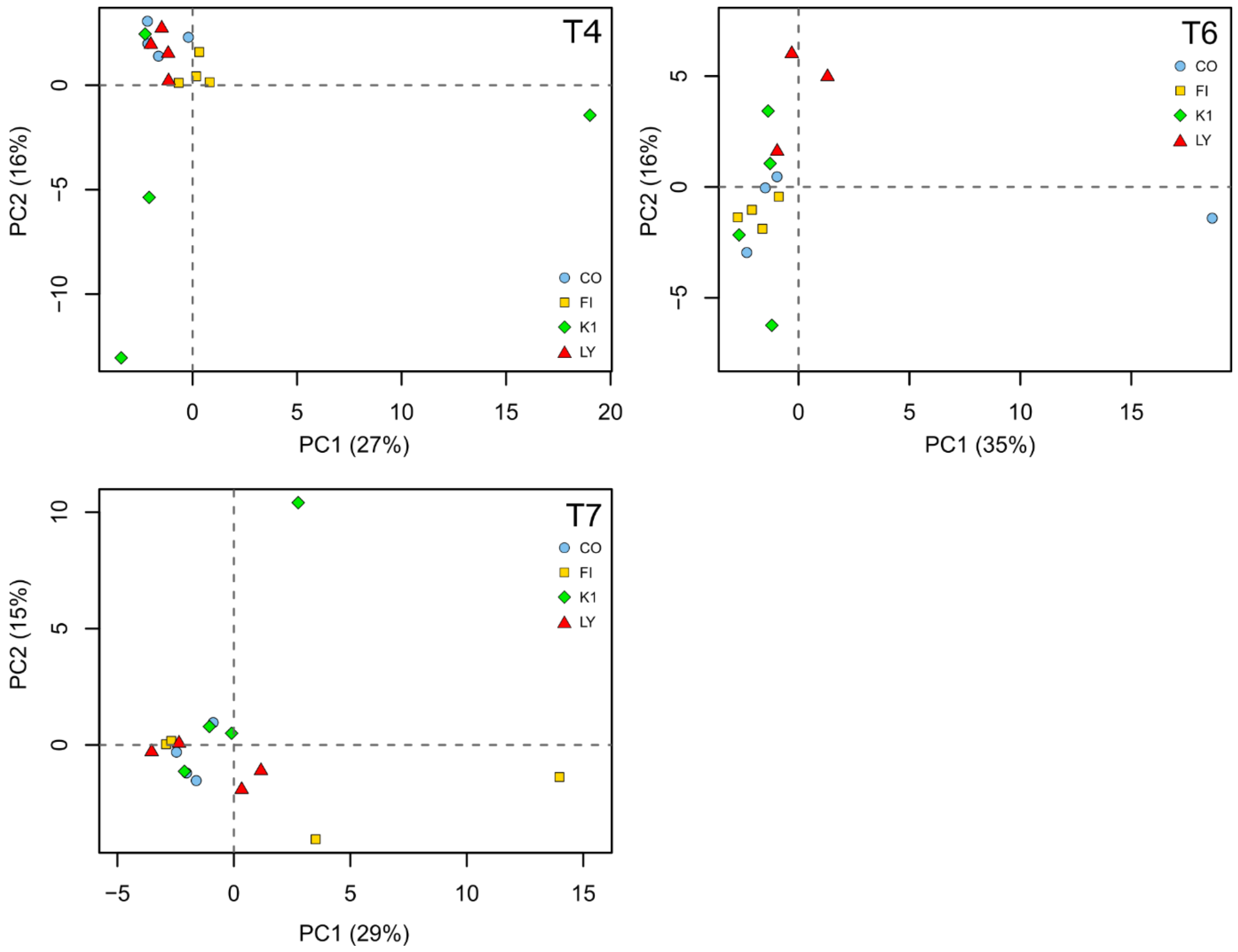


**Figure 10.** Principal component analysis plot of swab microbial relative abundances (gut) at each timepoint, and treatment: control (CO), filamentous phage (FI), *Vibrio* (K1) and lytic phage (LY).
